# Tuning of liver circadian transcriptome rhythms by thyroid hormone state in male mice

**DOI:** 10.1101/2023.09.23.559037

**Authors:** Leonardo Vinícius Monteiro de Assis, Lisbeth Harder, José Thalles Lacerda, Rex Parsons, Meike Kaehler, Ingolf Cascorbi, Inga Nagel, Oliver Rawashdeh, Jens Mittag, Henrik Oster

## Abstract

Thyroid hormones (THs) are important regulators of systemic energy metabolism. In the liver, they stimulate lipid and cholesterol turnover and increase systemic energy bioavailability. It is still unknown how the TH state interacts with the circadian clock, another important regulator of energy metabolism. We addressed this question using a mouse model of hypothyroidism and performed circadian analyses. Low TH levels decreased locomotor activity, food intake, and body temperature mostly in the active phase. Concurrently, liver transcriptome profiling showed only subtle effects compared to elevated TH conditions. Comparative circadian transcriptome profiling revealed alterations in mesor, amplitude, and phase of transcript levels in the livers of low-TH mice. Genes associated with cholesterol uptake, biosynthesis, and bile acid secretion showed reduced mesor. Increased and decreased cholesterol levels in the serum and liver were identified, respectively. Combining data from low- and high-TH conditions allowed the identification of 516 genes with mesor changes as molecular markers of the liver TH state. These genes participate in many known TH-associated processes. We further explored these genes and created a unique expression panel that can assess liver TH state in a time-of-day dependent manner. Our findings suggest that the liver has a low TH state under physiological conditions. Circadian profiling reveals genes as potential markers of liver TH state in one-time point studies.

## INTRODUCTION

Thyroid hormones (THs) are important regulators of embryonic development and energy metabolism. Produced in the thyroid gland in response to stimulation of the hypothalamus-pituitary-thyroid axis, thyroxine (T_4_) and, to a much lower extent, the biologically active 3,3’,5-triiodothyronine (T_3_) are secreted into the bloodstream. At target tissues, T_4_ is converted into T_3_ via specific deiodinases (DIOs). T_3_ can bind to the two nuclear TH receptors (THR) alpha/THRA and beta/THRB which act as transcription factors. As in most tissues, T_3_ action in the liver is predominately exerted by one primary nuclear receptor, in this case, THRB [1–3].

THs effects in mammals are diverse and highly tissue specific. The thyroid state (i.e., systemic levels THs) profoundly affects energy metabolism with high THs levels correlating with lower body weight and increased thermogenesis, lipolysis, and glucose usage. In the liver, T_3_ simultaneously induces *de-novo* lipid biosynthesis and, to a greater extent, lipolysis. Increased liver fatty acid (FA) uptake and turnover through beta-oxidation in mitochondria and peroxisomes are induced by T_3_. Similarly, T_3_ enhances cholesterol uptake, biosynthesis, and metabolization into bile acids, and it also increases hepatic gluconeogenesis and inhibits glycolysis and acetyl-coA utilization in the tricarboxylic acid (TCA) cycle [1,3,4]. Low THs levels, found in subclinical and clinical hypothyroidism, are associated with an increased incidence of non-alcoholic fatty liver disease (NAFLD) or Metabolic Dysfunction Associated Steatotic Liver Disease (MASLD) [5–8].

While the general effects of TH on liver metabolism are well-characterized, it remains largely unknown how the thyroid state interacts with the circadian regulation of physiological processes in an organ. Most species have developed endogenous time-keeping mechanisms that allow them to keep track of (day-) time and adjust physiology and behavior in anticipation of regularly recurring events. In mammals, a central clock residing in the hypothalamic suprachiasmatic nucleus (SCN) is reset by the external light-dark cycle and coordinates molecular oscillators in central and peripheral tissues including the liver. At the molecular level, a series of oscillatory interlocked transcriptional-translational feedback loops comprised of clock genes and proteins oscillate throughout the day, adjusting cellular functions across the 24-hour day cycle. Rhythmic factors such as body temperature, hormones (e.g., cortisol and melatonin), and autonomic nervous stimuli are known pathways through which the SCN pacemaker regulates peripheral clocks and rhythms [9,10], but it is still up for discussion how low amplitude rhythmic or arrhythmic signals interact with circadian functions at the tissue level.

We have recently shown that, in mice, a high-TH state leads to marked time-of-day specific alterations in energy metabolism such as increased energy expenditure during the active (i.e., the night) and higher body temperature during the inactive phase (i.e., the day). In the liver, T_3_ treatment leads to a rewiring of the diurnal liver transcriptome with strong effects on energy metabolism-associated genes, especially those involved in glucose and lipid metabolism. Transcriptional and metabolite data suggest higher triglyceride biosynthesis during the inactive phase followed by increased lipolysis rates in the active phase in T_3_-treated mice. Interestingly, these effects are independent of the liver clock gene machinery itself, which is rather insensitive to T_3_ treatment [11].

While the effects of high-T_3_ in the liver have been characterized, we investigated here how a low thyroid state affects the circadian regulation of energy metabolism and the liver transcriptome. Low TH levels reduced systemic energy turnover in line with human hypothyroid conditions. At the same time, only modest effects were observed on the liver transcriptome. Circadian transcriptome profiling allowed for more fine-grained characterization of low-TH effects revealing changes in lipid and cholesterol metabolism. Our approach identified several temporally stable TH state-responsive genes that may serve as livers-specific biomarkers of the TH state.

## MATERIAL AND METHODS

### Mouse model and experimental conditions

Two- to three-months-old male C57BL/6J mice (Janvier Labs, Germany) were housed in groups of three under a 12-hour light, 12-hour dark (LD, ∼300 lux) cycle at 22 ± 2 °C and a relative humidity of 60 ± 5 % with *ad-libitum* access to food and water. Mice were treated with methimazole (MMI, 0.1%, Sigma-Aldrich, USA) potassium perchlorate (0.2%) and sucralose (1 tablet per 50 ml, Tafelsüss, Borchers) for three weeks.

During the treatment period, mice were monitored for body weight individually and food and water intake per cage. All *in vivo* experiments were ethically approved by the Animal Health and Care Committee of the Government of Schleswig-Holstein and were performed according to international guidelines on the ethical use of animals. The sample size was calculated using G-power software (version 3.1) and are shown as biological replicates in all graphs.

Euthanasia was carried out using cervical dislocation and tissues were collected every 4 h. Night experiments were carried out under dim red light. Tissues were immediately placed on dry ice and stored at -80 °C until further processing. Blood samples were collected from the trunk, and clotting was allowed for 20 min at room temperature. Serum was obtained after centrifugation at 2,500 rpm, 30 min, 4 °C, and samples stored at -20 °C.

### Total T_3_ and T_4_ evaluation

Serum quantification of T_3_ and T_4_ was performed using commercially available kits (NovaTec, Leinfelden-Echterdingen, DNOV053, Germany for T_3_ and DRG Diagnostics, Marburg, EIA-1781, Germany for T_4_) following the manufactures’ instructions.

### Triglycerides and cholesterol evaluation

TAG and total cholesterol evaluation were processed according to the manufacturer’s instructions (Sigma-Aldrich, MAK266 for TAG and Cell Biolabs, San Diego, USA, STA 384 for cholesterol).

### Telemetry and metabolic evaluation

Core body temperature and locomotor activity were monitored in a subset of single-housed animals using wireless transponders (E-mitters, Starr Life Sciences, Oakmont, USA). Probes were transplanted into the abdominal cavity of mice 7 days before starting the drinking water treatment. During the treatment period, mice were recorded once per week for at least two consecutive days. Recordings were registered in 1-min intervals using the Vital View software (Starr Life Sciences). Temperature and activity data were averaged over two consecutive days (treatment days: 19/20) and plotted in 60-min bins.

An open-circuit indirect calorimetry system (TSE PhenoMaster, TSE Systems, TSE Systems, USA) was used to determine respiratory quotient (RQ = carbon dioxide produced / oxygen consumed) and energy expenditure in a subset of single-housed mice during drinking water treatment. Mice were acclimatized to the system for one week prior to starting the measurement. Monitoring of oxygen consumption, water intake as well as activity took place simultaneously in 20-min bins. VO_2_ and RQ profiles were averaged over two consecutive days (treatment days: 19/20) and plotted in 60-min bins. Energy expenditure was estimated by determining the caloric equivalent according to Heldmaier [12]: heat production (mW) = (4.44 + 1.43 * RQ) * VO_2_ (ml O_2_/h).

### Microarray analysis

Total RNA was extracted using TRIzol (Thermofisher, Waltham, USA) and the Direct-zol RNA Miniprep kit (Zymo Research, Irvine, USA) according to the manufacturer’s instructions. Genome-wide expression analyses was performed using Clariom S arrays (Thermo Fisher Scientific) using 100 ng RNA of each sample according to the manufacturer’s recommendations (WT Plus Kit, Thermo Fisher Scientific). Data were analyzed using Transcriptome Analyses Console (Thermo Fisher Scientific, version 4.0) and expressed in log_2_ values. Sample MMI_ZT06_a and MMI_ZT18_d were removed from the data due to low quality.

### Differentially expressed gene (DEG) analysis

To identify global DEGs, all temporal data from each group was considered and analyzed by *Student’s t* test and corrected for false discovery rates (FDR < 0.1). Up- or downregulated DEGs were considered when a threshold of 1.5-fold (0.58 in log_2_ values) regulation was met. As multiple probes can target a single gene, we curated the data to remove ambiguous genes. To identify DEGs at specific time points (ZTs – Zeitgeber time; ZT0 = “lights on”), the procedure described above for each ZT was performed separately. Time-independent DEGs were identified by finding consistent gene expression pattern across all ZTs.

### Rhythm analysis

To identify probes that showed diurnal (i.e., 24-hour) oscillations, we employed the non-parametric JTK_CYCLE algorithm [13] in the Metacycle package [14] with a set period of 24 h and an adjusted p-value (ADJ.P) cut-off of 0.05. Phase and amplitude parameter estimates from CircaSingle were used for rose plot visualizations [15]. To directly compare rhythm parameters (mesor and amplitude) in gene expression profiles between T_3_ and CON, CircaCompare fits were used irrespective of rhythmicity thresholds. Phase comparisons were only performed when a gene was considered as rhythmic in both conditions (p < 0.05) as previously described [11]. Temporal profiles were made using geom_smooth (ggplot2 package), method “lm”, and formula = y ∼ 1 + sin(2*pi*x/24) + cos(2*pi*x/24). Small differences in rhythmic parameters can be present due to the different sine curve fitting between CircaCompare and ggplot curve fit.

### Gene set enrichment analysis (GSEA)

Functional enrichment analysis of DEGs was performed using the Gene Ontology (GO) annotations for Biological Processes on the Database for Annotation, Visualization, and Integrated Discovery software (DAVID 6.8 [16]). Processes were considered significant for a biological process containing at least 5 genes (gene count) and a p-value < 0.05. To remove the redundancy of GSEA, we applied the REVIGO algorithm [17] using default conditions and a reduction of 0.5. For enrichment analyses from gene sets containing less than 100 genes, biological processes containing at least 2 genes were included. Overall gene expression evaluation of a given biological process was performed by normalizing each timepoint by CON mesor.

### Data handling and statical analysis of non-bioinformatic related experiments

Samples were only excluded upon technical failure. Data from ZT0-12 were considered as light phase and from ZT 12 to 24 as dark phase. Day *vs.* night analyzes were performed by averaging the data and comparing using unpaired *Student’s* t test with Welch correction. Time-course data were analyzed by Two-way ANOVA for main treatment effects followed by Bonferroni post-test. When applicable, Two-Way for repeated measures was applied. Single timepoint data were evaluated by unpaired *Student’s* t test with Welch correction or Mann-Whitney test for parametric or non-parametric samples, respectively. ANOVA One-Way followed by Tukey was used for single timepoint data when CON, MMI, and T_3_ groups were evaluated. Spearman’s correlation was used for all correlational analyzes. Analyzes were done in Prism 9.4 (GraphPad) and a p-value < 0.05 was used to reject the null hypothesis.

### Data handling and statical analysis of bioinformatic experiments

Statistical analyses were conducted using R 4.0.3 (R Foundation for Statistical Computing, Austria) or in Prism 9.4 (GraphPad). Rhythmicity was calculated using the JTK_CYLCE algorithm in meta2d, a function of the MetaCycle R package v.1.2.0. Rhythmic features were calculated and compared pairwise among the groups using the CircaCompare R package v.0.1.1. Data visualization was performed using the ggplot2 R package v.3.3.5, eulerr R package v.6.1.1, UpSetR v. 1.4.0, pheatmap v. 1.0.12, and Prism 9.4 (GraphPad).

### Data mining from published studies

Data from CON and T_3_ were used and properly disclosed in each section. All experimental data from [11] was extracted from the <u>Figshare</u> depository, unless specified.

Microarray data was extracted from Gene Expression Omnibus (GEO) database (GSE199998).

### Data availability

All experimental data are already deposited in public depositories. No additional data was generated.

## RESULTS

### TH state tunes systemic energy metabolism

To model hypothyroid conditions in mice, we supplemented their drinking water with methimazole and potassium perchlorate (MMI group) which suppresses TH biosynthesis through inhibition of thyroperoxidase in the thyroid gland and inhibits iodine uptake [18]. MMI treatment resulted in a strong reduction of T_3_ and T_4_ levels. T_3_ levels showed circadian rhythms in the control (CON) and MMI groups peaking in the late and mid-dark phase, respectively (Figure 1A). T_4_, on the other hand, was arrhythmic in both conditions (Figure 1B; Supplementary file 1).

**Figure 1:**
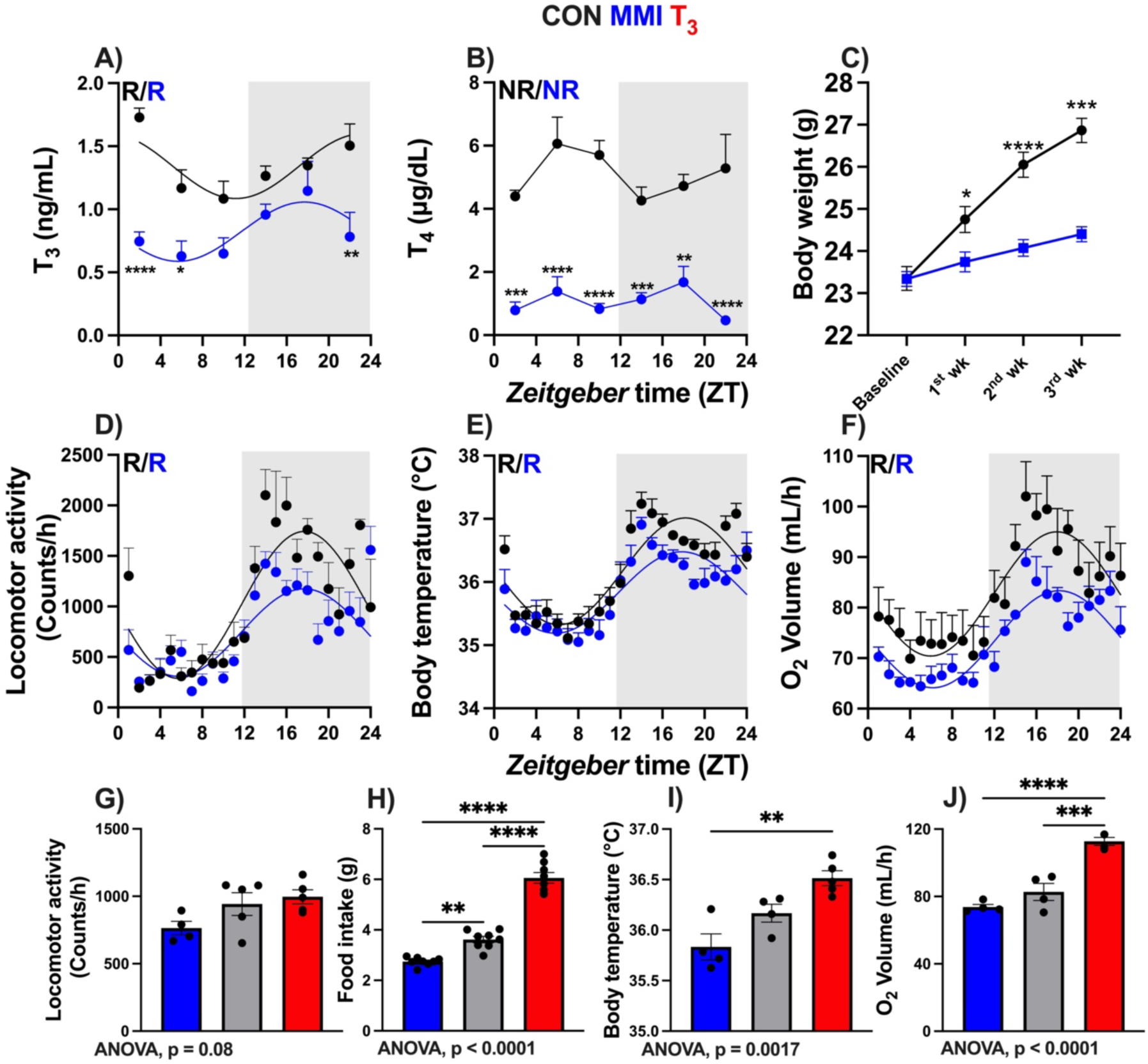
Dose-dependent effects of thyroid hormone (TH) state on systemic energy metabolism. A – F) Serum levels of T_3_ and T_4_, body weight across the experiment, 24-hour profiles of locomotor activity, body temperature, and O_2_ consumption are shown. Rhythmicity was assessed using CircaCompare algorithm. Presence (R) or absence (NR) of significant circadian rhythmicity is depicted. In the presence of significant 24-hour rhythmicity, a sine curve was fit to the data. G – J) 24h average data for locomotor activity, food intake, body temperature, and O_2_ consumption for MMI, CON, and T_3_ (de Assis et al., 2022) groups. One- way ANOVA was performed (p value is shown) followed by Tukey’s post-test comparisons (depicted with asterisks). In A and B, n = 4 – 6 animals per group or timepoint. In C, n = 24. In D, n = 4 and 5 for CON and MMI groups, respectively. In E and F, n = 4 for each group. *, **, ***, **** represent a p value of < 0.05, 0.01, 0.001, and 0.0001, respectively.

Reduced food and water intake were observed in MMI compared to CON mice (Supplementary figure 1A – B). As had been reported before [19] – and unlike hypothyroid humans – MMI mice had reduced body weight compared to CON mice (Figure 1C). Compared to CON mice, locomotor activity rhythms were similar in MMI mice, but activity was reduced specifically during the dark phase (Figure 1D; Figure 1 – Figure Supplement 1C). Body temperature was reduced in MMI during the dark phase (Figure 1E; Supplementary figure 1D). Energy turnover, assessed by oxygen consumption, was slightly lower throughout the day (Figure 1F; Supplementary figure 1E) while respiratory quotient profiles were largely unaltered in MMI mice (Figure 1; Supplementary figure 1 F – G).

By averaging profile data over the whole day and comparing data to high-T_3_ conditions [11], clear systemic effects of TH state on metabolic homeostasis became apparent. TH state-dependent effects were observed for locomotor activity (r = 0.62, p = 0.0191), food intake (r = 0.85, p = 0.0002), body temperature (r = 0.75, p = 0.0042), and oxygen consumption (r = 0.79, p = 0.032, Figure 1 G – J). In sum, our findings confirm that a low TH state decreases systemic energy turnover, but this effect is more pronounced during the dark (active) phase.

### Low TH levels have moderate effects on liver transcription

In face of the marked effects of TH state on systemic energy metabolism, we focused our attention on the liver due to its major role as a metabolic organ. We collected tissues every 4h over the whole day to allow the evaluation of time-of-day dependent effects of TH state on liver transcription. To assess T_3_ state effects, we determined differentially expressed genes (DEGs) in T_3_ and MMI conditions irrespective of sampling time. Surprisingly, T_3_ mice showed a 5-fold higher number of DEGs compared to MMI mice (Figure 2A; Supplementary File 2). DEGs analysis performed separately for each sampling time yielded 95 DEGs in MMI livers compared to 2,200 DEGs (a factor of ca. 23) in high-T_3_ mice [11] (Figure 2B – C; Supplementary File 2). No robust DEGs, i.e., genes consistently differentially expressed across all sampling times (ZTs), were found in MMI livers (compared to 37 robust DEGs in T_3_ treated mice [11] (Figure 2 D). Analysis of established liver TH/THR target genes and modulators confirmed a much stronger effect of high-T_3_ than low TH conditions on liver TH state (Figure 2 E; Supplementary figure 2).

**Figure 2:**
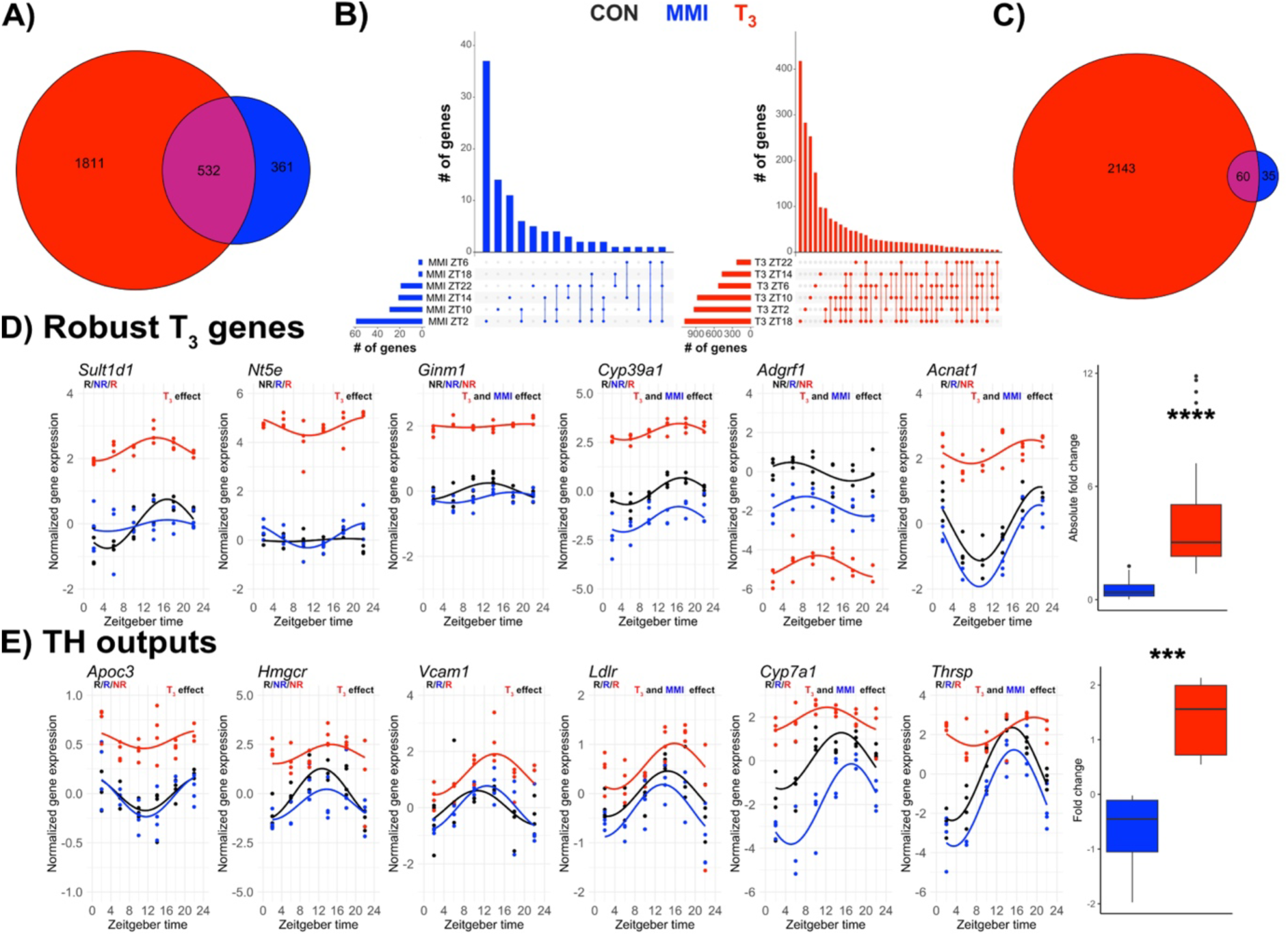
Lowering thyroid hormone state has subtle effects on liver transcriptome rhythms. A) Global DEG analysis (disregarding sampling time) is represented as a Venn diagram. B) Upset plots represent DEG analysis for each ZT separately. C) Venn diagram represents all temporal DEGs (i.e., showing different expression levels of at least one ZT) identified in MMI and T_3_ groups versus CON mice. D) Selected examples of robust DEGs previously identified in T_3_-treated mice. Absolute fold change comparison of all 37 DEGs in T_3_ and MMI mice are shown. E) Selective TH output genes and fold change of these genes. Comparisons were performed using two-way ANOVA (main treatment effect, p < 0.05). n = 3 – 4 for all ZTs and groups. Pair-wise comparisons were performed by Student’s t test with Welch correction. Presence (R) or absence (NR) of circadian rhythm by JTK cycle (p value < 0.05). ***, **** represents a p value of < 0.001, and 0.0001, respectively.

Together, these data show that despite marked systemic effects, MMI treatment had much lower effects on liver transcription. Vice versa, this suggests that, under physiological conditions (CON), the liver is already in a low-TH state.

### Diurnal profiling reveals transcriptional effects of a low-TH state in the liver

Evaluation of core clock gene expression profiles showed little effect on rhythmic parameters between MMI and CON mice (Figure 3A) in line with what was previously observed under high-T_3_ conditions [11]. To fully assess the rhythmic liver transcriptome, the JTK cycle algorithm [13] was used. A total of 3,329 and 3,383 genes (3,354 and 3,397 probe sets) were classified as rhythmic (p < 0.05) in CON and MMI, respectively (Supplementary figure 3). 1,412 genes (1,417 probes) were significantly rhythmic in both groups (Supplementary figure 3). An average phase delay of 0.24h was observed for these genes in MMI livers (Figure 3 B) compared to a 1h phase advance in high-T_3_ conditions [11]. Expression peak phases of rhythmic genes were widely distributed, and amplitudes were overall similar between conditions (Supplementary figure 3).

**Figure 3:**
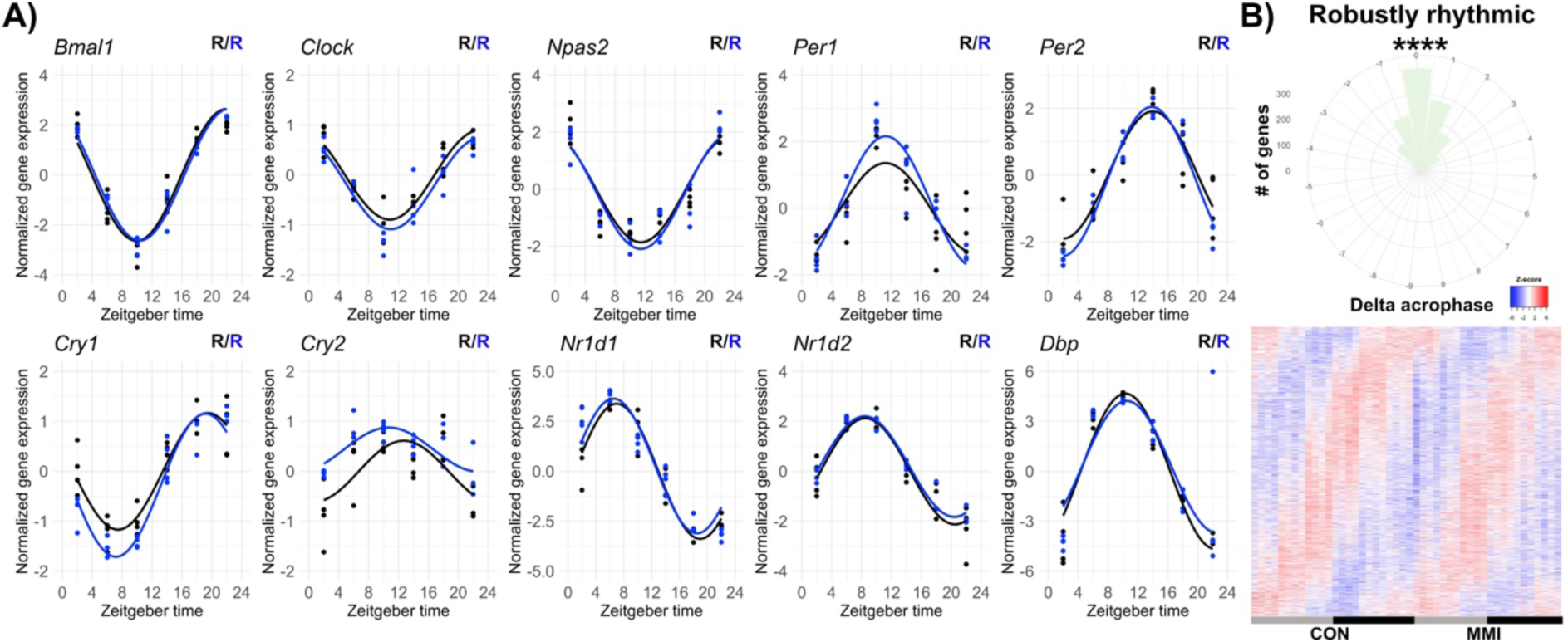
Lowering thyroid hormone state does not affect the rhythmic expression of core clock genes and has a slight phase effect on robustly rhythmic genes. A) Diurnal expression profile of core clock genes is shown. Presence (R) or absence (NR) of significant circadian rhythm by JTK cycle (p value < 0.05) is depicted. B) Rose plot and heatmap of robustly rhythmic genes are shown. A minor phase delay of 0.24 h was identified in the MMI group compared to CON (test against zero, p < 0.001). n = 3 – 4 for all ZTs and groups.

To investigate the difference in rhythmic characteristics of the transcriptome regulation under low TH conditions, we performed differential circadian rhythm analysis using CircaCompare [15]. Of the 5,297 genes (5,334 probes) showing significant 24h rhythmicity in at least one group, 1,882 genes showed changes in mesor, 403 in amplitude, and 391 in phase between low-TH and control conditions (Figure 4 A – B; Supplementary file 4). Processes associated with cellular proliferation, mRNA processing, and macromolecule complex assembly were enriched in genes with mesor DOWN while processes associated with response to oxidative stress and xenobiotic metabolism processes were found in the mesor UP genes. Genes associated with response to hypoxia, exocytosis, and lipid transport showed increased amplitudes in MMI livers. Conversely, the expression of genes involved in oxidative stress and detoxification processes (cellular oxidant detoxification, hydrogen peroxide catabolism, glutathione, and ethanol metabolism) were phase-delayed in MMI mice (Figure 4 C; Supplementary file 4).

**Figure 4:**
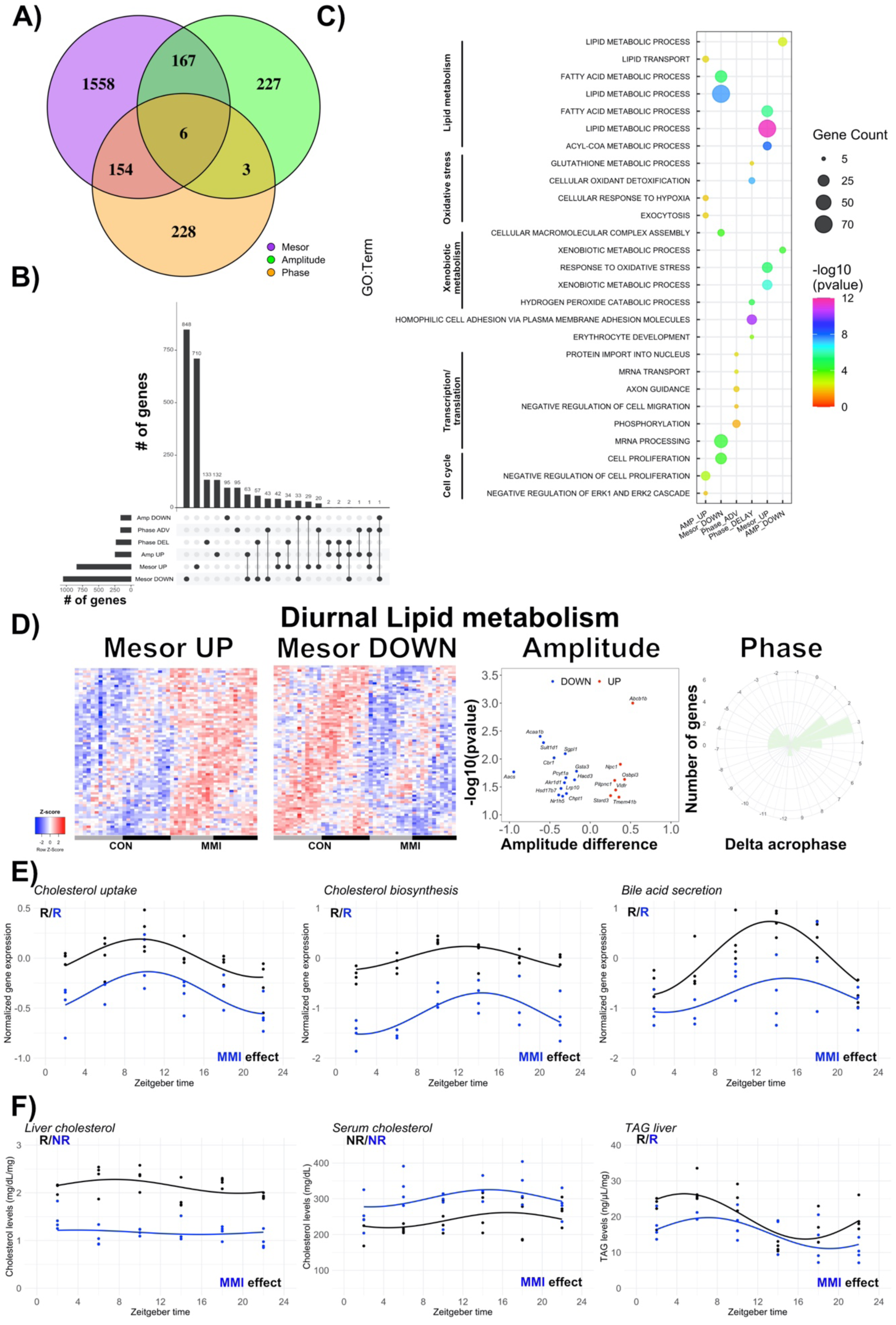
Differential rhythm analysis reveals changes in liver transcriptome rhythms that affect lipid and cholesterol metabolism in MMI mice. A) Differential rhythm analysis was performed using CircaCompare and is represented as Venn diagrams. B) Upset plots show alterations in diurnal rhythm parameters (mesor, amplitude, phase). C) Gene set enrichment analysis (GSEA) of the genes with either mesor, amplitude, or phase alterations was performed. Top-5 biological processes for each category are shown. D) In-depth diurnal lipid metabolism analysis in response to low thyroid hormone state. Heatmaps show genes with mesor changes. Volcano plot and rose plot show alterations in amplitude and phase, respectively. E) Normalized gene expression of selected genes participating in cholesterol uptake, biosynthesis, and degradation (bile acid secretion) in CON and MMI mice. F) Quantification of cholesterol and TAG in serum or liver. Presence (R) or absence (NR) of significant circadian rhythm by CircaCompare (p value < 0.05) is depicted. n = 3 – 4 for all ZTs and groups.

Lipid metabolism-associated processes were regulated for all three rhythm parameters (Figure 4 C). A dual effect (up- and down-regulation) on mesor was observed while most lipid metabolism genes showed reduced amplitudes and phase delays in MMI mice (Figure 4 D). The identified genes were manually inspected and only those genes participating directly in lipid metabolism pathways were included. In this refined gene set for mesor UP, approximately 50% of the genes were associated with acyl-CoA degradation (*Acot1*, *2*, *3*, *4*, *7*, *9*, *13*) and biosynthesis (*Acsm 1*, *3*, *5*) while others were associated with lipolysis (*Lpl*, *Lipa*) and TAG uptake (*Vldr*). In the mesor DOWN group, ca. 40% of the genes were involved in cholesterol biosynthesis (*Aacs*, *Dhcr7*, *Hmgcr*), uptake (*Ldlr*, *Lrp10*, *Npc1*, *Lrp5*, *Pcsk9*), and bile acid secretion (*Abcb11*, *Cyp7a1*, *Abcg5*, *Slc10a1*, *Slc10a5*, *Slc10a2*). Moreover, 35% of the genes were associated with FA biosynthesis (*Acacb*, *Fasn*), elongation (*Elovl1*, *2*, *3*, *5*, *Hacd1*), transport (*Cpt1a*, *Fabp5*), and uptake (*Slc27a2*, *Slc27a5*). Around 30% of amplitude DOWN genes were associated with cholesterol biosynthesis (*Aacs*, *Hsd17b7*) and secretion (*Akr1d1*, *Nr1h5*) while genes with a gain in amplitude were linked with cholesterol uptake (*Abcb1b*) and internal trafficking (*Stard3*, *Npc1*) as well as TAG uptake (*Vldlr*). Genes with expression profile phase effects were more diverse in their functions. Phase-delayed genes were associated with cholesterol metabolization (*Cyp8b1*) and bile acid secretion (*Cyp7a1*, *Abcb11*), FA biosynthesis (*Fads2*), transport (*Cpt1a*) and elongation (*Elovl2*) (Supplementary file 4).

Focusing on cholesterol metabolism, we divided this pathway into three categories: uptake, biosynthesis, and bile acid secretion. Averaged diurnal gene expression from both groups was rhythmic with a strong mesor DOWN effect in MMI mice (Figure 4E). Total liver cholesterol was reduced and arrhythmic in MMI mice compared to the CON group. On the other hand, serum total cholesterol was elevated in MMI compared to CON mice, indicating impairment in liver cholesterol uptake – as previously suggested. Liver TAG was less responsive to a low-TH state and showed a mesor DOWN effect (Figure 4F).

Our findings show that a low-TH state results in distinct alterations in liver transcriptome rhythms. Rhythms in lipid and cholesterol metabolism-associated gene programs are affected by low TH levels and translate into differences mostly in cholesterol levels.

### Identification of liver TH response genes

When assessing liver TH state across studies – particularly those based in clinical settings – it is not always possible to consider temporal dynamics in liver physiology. Therefore, it would be helpful to identify molecular markers that indicate liver TH state largely independent of sampling time. However, unlike what we had previously shown for high-TH conditions, no robust DEGs were identified in livers of MMI mice (Figure 2B). While this would preclude reliable data analysis independent of sampling time, we circumvented this limitation by filtering our dataset for transcripts with marked TH state-dependent mesor effects in expression. This would allow for reliable detection of TH state at any time point if sampling time were consistent across experimental conditions. We identified 516 genes that showed consistent mesor effects in response to changes in TH state for both MMI and T_3_ conditions (Figure 5 A). Gene Set Enrichment Analysis (GSEA) for genes in the DOWN-MMI/UP-T_3_ group showed enrichment for metabolic processes such as steroid, cholesterol, FA, and bile acid metabolism. Conversely, genes in the UP-MMI/DOWN-T_3_ group were associated with processes involved in FA, lipoprotein, carbohydrate, and xenobiotic metabolism, amongst others. Interestingly, glucose Glucose/glycogen metabolism was exclusively enriched in the UP in MMI/down in T_3_ group (Figure 5 B – C; Supplementary file 5), suggesting a TH-driven overall inhibition of these pathways in line with our previous observations [11]. Averaged expression of all genes pertaining to similar biological processes across the day showed that lipid metabolism – mostly comprised of FA biosynthesis, TAG uptake and biosynthesis – genes were highly responsive to TH levels, being up- and downregulated in T_3_ and MMI mice, respectively. Genes associated with FA catabolism were up- and downregulated in MMI and T_3_ mice, respectively. As already indicated in the GSEA, cholesterol and bile acid metabolism were markedly affected as MMI and T_3_ mice showed down- and upregulation, respectively (Figure 5 C; Supplementary file 5).

**Figure 5:**
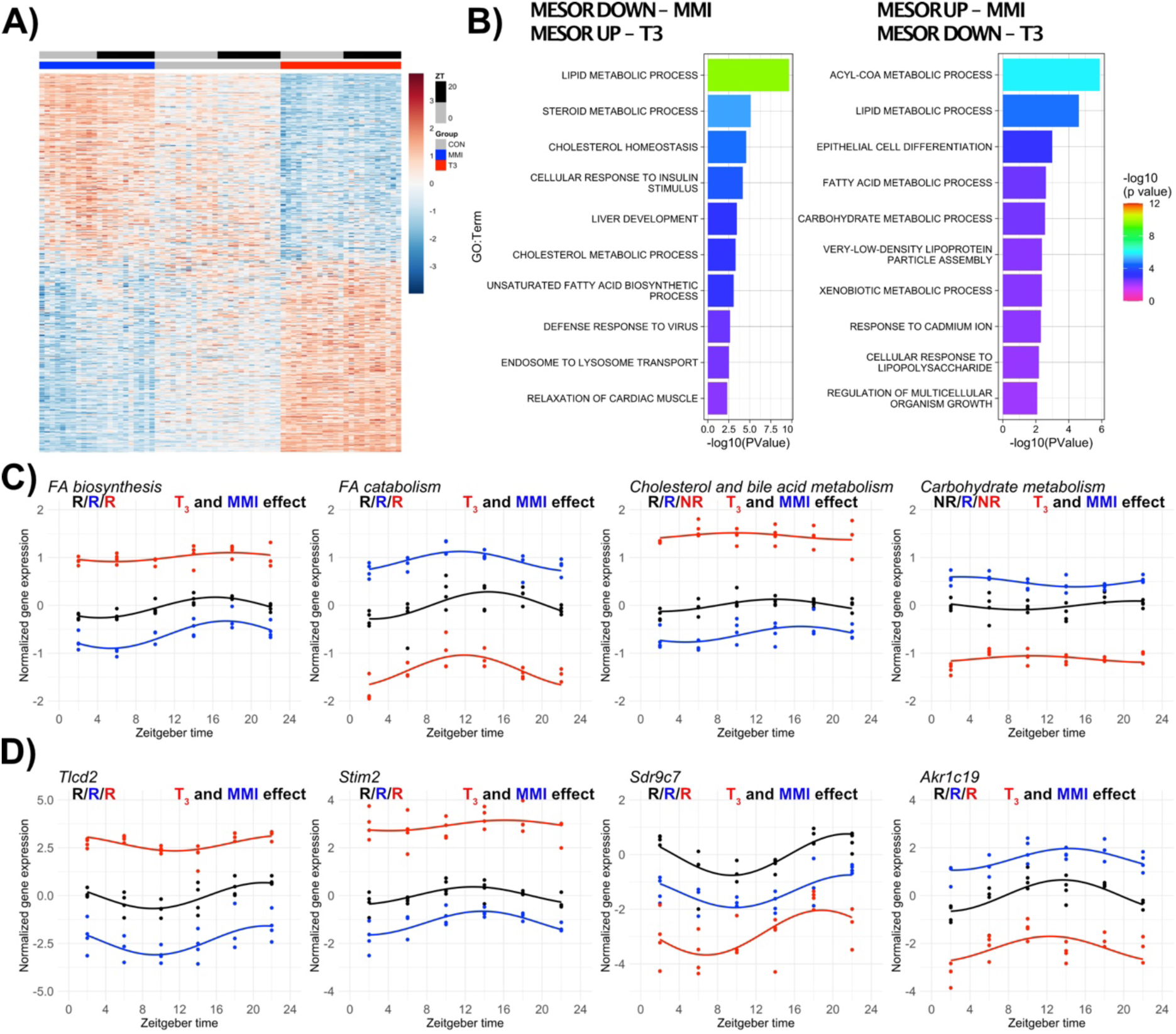
Identification of thyroid hormone-responsive genes by mesor comparison. A) Heatmap shows the 516 TH-dose responsive genes with differential mesor expression among each group. B) Gene set enrichment analysis (GSEA) of mesor-altered genes are shown for each condition. C) Genes pertaining to similar biological pathways identified in B were normalized by CON mesor and plotted. FA biosynthesis pathway comprises lipid metabolic process, unsaturated fatty acid biosynthetic process, long-chain fatty-acyl-CoA biosynthetic process, diacylglycerol biosynthetic process, negative regulation of fatty acid biosynthetic process, and lipid storage. FA catabolism pathway is comprised of acyl-CoA metabolic process, lipid metabolic process, fatty acid metabolic process, very-low-density lipoprotein particle assembly processes. Cholesterol and bile acid metabolism pathways comprise cholesterol homeostasis, cholesterol metabolic process, steroid metabolism, and bile acid signaling pathway. Carbohydrate metabolism pathways are comprised of carbohydrate metabolic process, glycogen metabolic process, ATP metabolic process, and glucose homeostasis. D) Selected biomarkers for TH state at all time points (additional genes can be found in Supplementary file 5). Presence (R) or absence (NR) of circadian rhythm by CircaCompare (p value < 0.05) is depicted. n = 3 – 4 for all ZTs and groups.

Absolute mesor change was used to rank each gene for each comparison (CON *vs.* T3 and CON *vs.* MMI). The top 4 genes (*Tlcd2*, *Stim2*, *Sdr9c7*, *Akr1c19*) are shown (Figure 5D) and recapitulate the low- and high- TH state at any fixed timepoint across the day. The fact that these genes are robustly rhythmic across conditions further emphasizes that sampling time should be kept consistent even in one-timepoint sampling studies (Figure 5 D; Supplementary file 5). To test the predictive efficiency of our approach, we used the published T_3_ atlas that combines transcriptome and TH receptor ChipSeq data from different tissues [20]. We identified an overlap of ca. 70% (179 of 251 genes) between the two datasets, further supporting the validity of our approach (Supplementary file 5).

## DISCUSSION

In line with the role of TH in metabolic regulation, reducing the TH state by MMI treatment decreased energy turnover in mice. Contrasting these systemic effects, transcriptional responses in the liver were surprisingly subtle with only a few temporally stable DEGs compared to animals with normal or elevated TH levels. Temporal profiling of transcriptome responses, however, revealed time-of-day dependent changes in metabolic pathways in MMI livers. An altered gene signature associated with TAG and cholesterol metabolism was identified. Interestingly, cholesterol levels were more sensitive to a low TH state than TAGs. Compiling our circadian low- and high-TH datasets genes, several biomarker genes for the TH state were identified and could be used to assess the liver TH state robustly.

In MMI mice, T_3_ and T_4_ levels were markedly suppressed, which confirms the success of our experimental approach. The circadian rhythm of T_3_ in the blood was retained at low-TH conditions. In line with previous studies in rodents and observations in hypothyroid patients, we identified TH dose- and time-dependent effects on systemic energy metabolism [19,21,22]. Notably, the effects of a low-TH state on energy metabolism were largely restricted to the active phase. MMI mice showed decreased body weight throughout the experiment, contrasting with hypothyroid conditions in humans. These differences were recently confirmed in genetic hypothyroidism mouse models and are associated with food intake suppression, higher skeletal-muscle adaptive thermogenic and fatty acid oxidation in low-TH mice [19].

Liver transcriptome analyses showed 23-fold fewer DEGs in MMI- compared to T_3_- treated (high-TH) mice. This reduced liver responsiveness to a low TH input is further reflected in decreased effects on the established TH/THR target genes [2,4]. Further, THR (*Thra* and *Thrb*) expression in response to TH state alterations was only seen in T_3_- but not in MMI-treated animals. In contrast, the expression of *Dio1*, the main deiodinase in the liver [23], showed a clear TH dose-dependent effect. Knockdown of liver *Dio1* favored the development of fatty liver disease (NAFLD/MASLD) due to reduced fatty acid oxidation [24]. Our data suggest that reducing TH action under physiological conditions would be of little consequence as the liver is already a low-TH organ. An alternative explanation for our findings of low liver responses to reduced TH state would be that intra-liver TH levels could be stabilized due to high DIO1 activity. *Dio1* mRNA was downregulated in the MMI group, which speaks against this idea, but evaluating DIO1 activity or directly measuring liver TH levels would be needed to exclude this option.

While these conclusions arise from a simple overall gene expression analysis, circadian profiling of transcriptome data allows for a much more fine-grained evaluation [25]. Previously undetected alterations in several genes revealed novel affected biological processes in response to a low TH state such as cellular proliferation, hypoxia, and oxidative stress, which may serve as hypothesis-generating data for follow-up experiments. We here focused our efforts on lipid metabolism because of the putative role of TH in NAFLD/MASLD regulation [26,27], a condition that also has been linked to circadian rhythms [28,29].

Gene signatures of lipid and cholesterol metabolism showed alterations in all three rhythm parameters (mesor, phase, and amplitude) in MMI-treated mice. Serum cholesterol was highly sensitive to TH state. Increased serum cholesterol levels in MMI mice agreed with previous observations of murine models for sub-clinical [30] or severe hypothyroidism [31]. Diurnal transcriptome profiling showed a mesor DOWN effect in genes associated with cholesterol uptake, biosynthesis, and bile acid secretion. A low TH state is known to reduce bile acid secretion in gall bladder, ileum, and feces compared to euthyroid mice [32] and an increased incidence of cholesterol gallstone formation has been associated with decreased circulating TH levels [33,34]. In contrast, liver TAG levels were less responsive to a low TH state despite marked related alterations at the transcript level. A murine study using a severe hypothyroidism model (low-iodine plus *Slc5a5* knockout) found a protective effect against NAFLD/MASLD. In this model, the hampered TH signaling impairs the adrenergic adipose-derived lipolysis, which reduces fatty acids shuttling to the liver. The reduction of fatty acid uptake by the liver prevents NAFLD/MASLD development. However, a mild hypothyroidism model, achieved by a low iodine diet for 12 weeks, increased liver TAG levels as the adrenergic adipose-derived fatty acid shuttling was not impaired [31]. Increased liver TAG has been reported in a subclinical model of hypothyroidism, achieved by low doses of MMI in drinking water after 16 weeks [30]. However, a recent study using propylthiouracil for 2 and 4 weeks found no changes in TAG and cholesterol levels despite changes in the serum [35].

Our findings show that a low TH state produces transcriptional changes that impair cholesterol metabolization by reducing liver cholesterol uptake, biosynthesis, and bile acid secretion. We suggest that the low cholesterol clearance of MMI mice results in increased serum cholesterol that can be classified as dyslipidemia and an early consequence of a low TH state. Although transcriptional changes associated with TAG metabolism were observed, liver TAG showed a less pronounced alteration and is suggestive of being a late consequence of hypothyroidism.

Considering that liver metabolism changes over the course of the day [36,37] and many metabolic genes showed time-of-day dependent responses to low- (this study) or high- [11] TH state, time emerges as an important variable in characterizing metabolic footprints in liver and, potentially, other tissues. We have recently modeled the impact of accounting for time in identifying DEGs in a mouse model of NASH motivated by low consistency in DEG identification in published studies. Accounting for time and using circadian analytical methods led to a ca. 7-fold higher DEG yield [25]

Circadian profiling may often not often be feasible – or simply too expensive – particularly in clinical settings. Therefore, we used our data to identify genes that consistently respond to changes in TH state at all times of day as potential robust markers of liver TH state. None of the previously identified 37 high-TH response genes [11] showed altered expression after MMI treatment. In this new approach, time was fully considered yielding markers of TH state in the liver. Rhythmic genes with mesor changes are associated with several metabolic processes known to be TH regulated such as lipid, glucose, steroid, and xenobiotic metabolism. We further explored the mesor-affected genes as possible candidates to estimate liver TH status. We restricted our analysis to genes that only showed mesor changes to avoid possible confounding factors caused by amplitude and/or phase. In this subset, 251 genes were identified as only being affected at the mesor level. We validated our approach in the recently published T_3_ atlas [20]. Comparing our mesor-restricted dataset (251 genes), we identified 179 DEGs that show a high overlap. Our findings show that several thousand T_3_-induced DEGs are affected by time, which can directly affect DEG detection if sampling time is inconsistent. However, our findings further refine this dataset and provide a selective set of genes that can be used to establish liver TH state. We suggest that this gene panel can be used to estimate liver TH state if sampling is time-controlled in experimental and possibly clinical studies.

As previously observed in a high-TH condition [11], the circadian core clock was largely unaffected by either low or high T_3_ conditions. These findings suggest that TH effects on transcriptome rhythms are downstream of the liver circadian clock and warrant further investigation. Metabolic changes evoked by a low TH state emerged only upon temporal profiling. We also identified a subset of genes that respond in a TH dose-dependent manner with changes in expression mesor. We suggest these genes can be used for follow-up validations of TH liver state in experimental conditions containing limited time points. Understanding the circadian effects of TH state may prove useful for identifying therapeutic targets and optimizing existing treatment strategies for metabolic liver diseases.

## CONFLICT OF INTEREST

All authors declare no competing interests that could have an impact on the study.

## ACKNOWLEDGEMENTS AND FUNDING

This work was supported by grants of the German Research Foundation (DFG) to HO 353-10/1, GRK-1957, and CRC/TR 296 “LOCOTACT” (ID 424957847, TP13 and TP14). JTH is a fellow of the São Paulo Research Foundation (FAPESP - 04524-8/2020).

---

**Supplementary figure 1:**
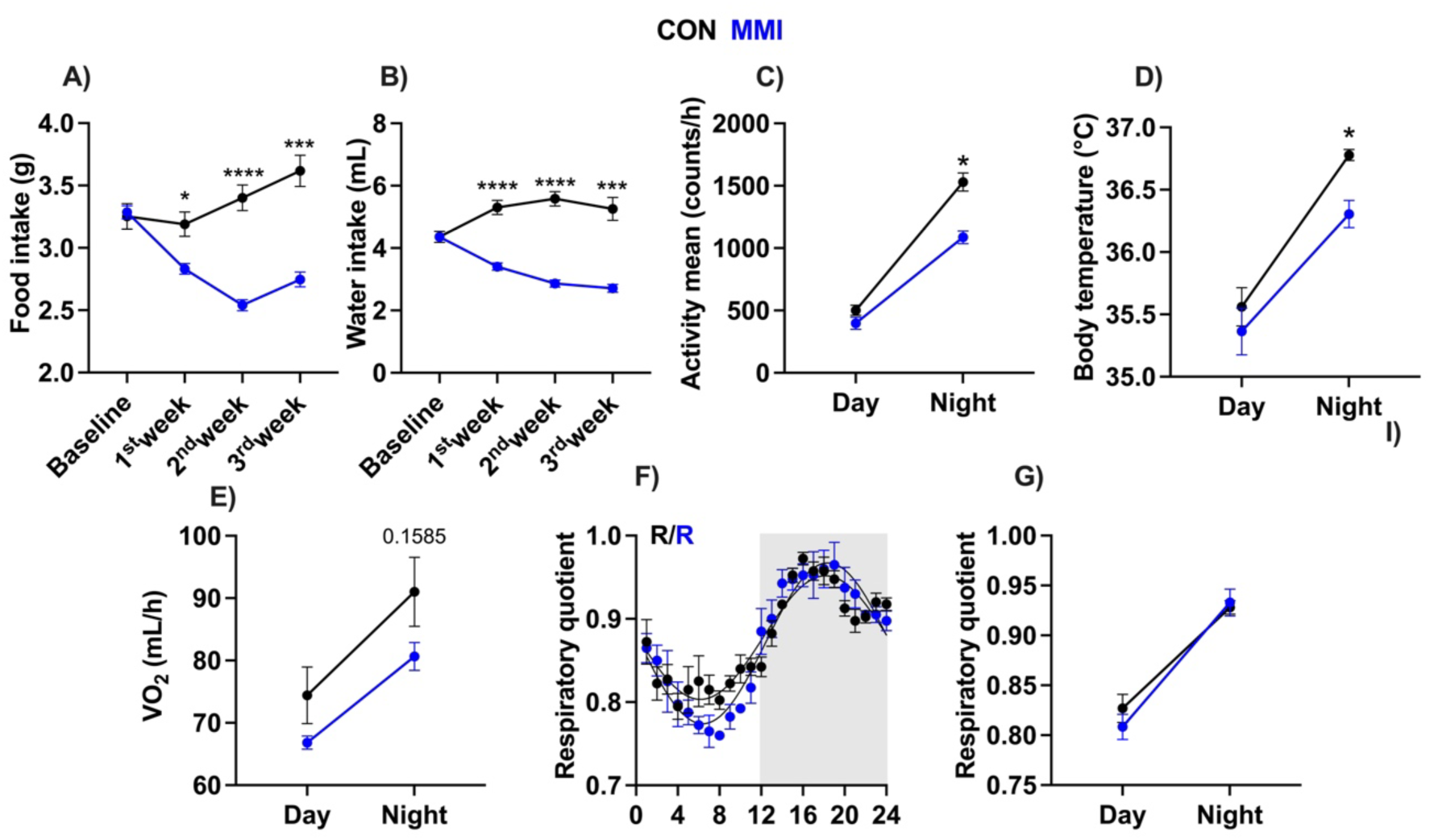
Evaluation of systemic metabolic parameters of CON and MMI mice. A – B) Assessment of food and water intake. C– E) Metabolic parameters (described in the y-axis) were obtained from the 3^rd^ week of the experiment. Day and night data were plotted by averaging values from ZT 0 to 12 (day) and from ZT 12 to 24 (night). Asterisks depict significant differences between CON and MMI mice. F – G) 24-hour profiles of respiratory quotient and day and night comparisons. In A and B, n = 8 (per cage). In C, n = 4 and 5 for CON and MMI groups, respectively. In D – G, n = 4 for each group. *, ***, **** represent a p value of < 0.05, 0.001, and 0.0001, respectively.

**Supplementary figure 2:**
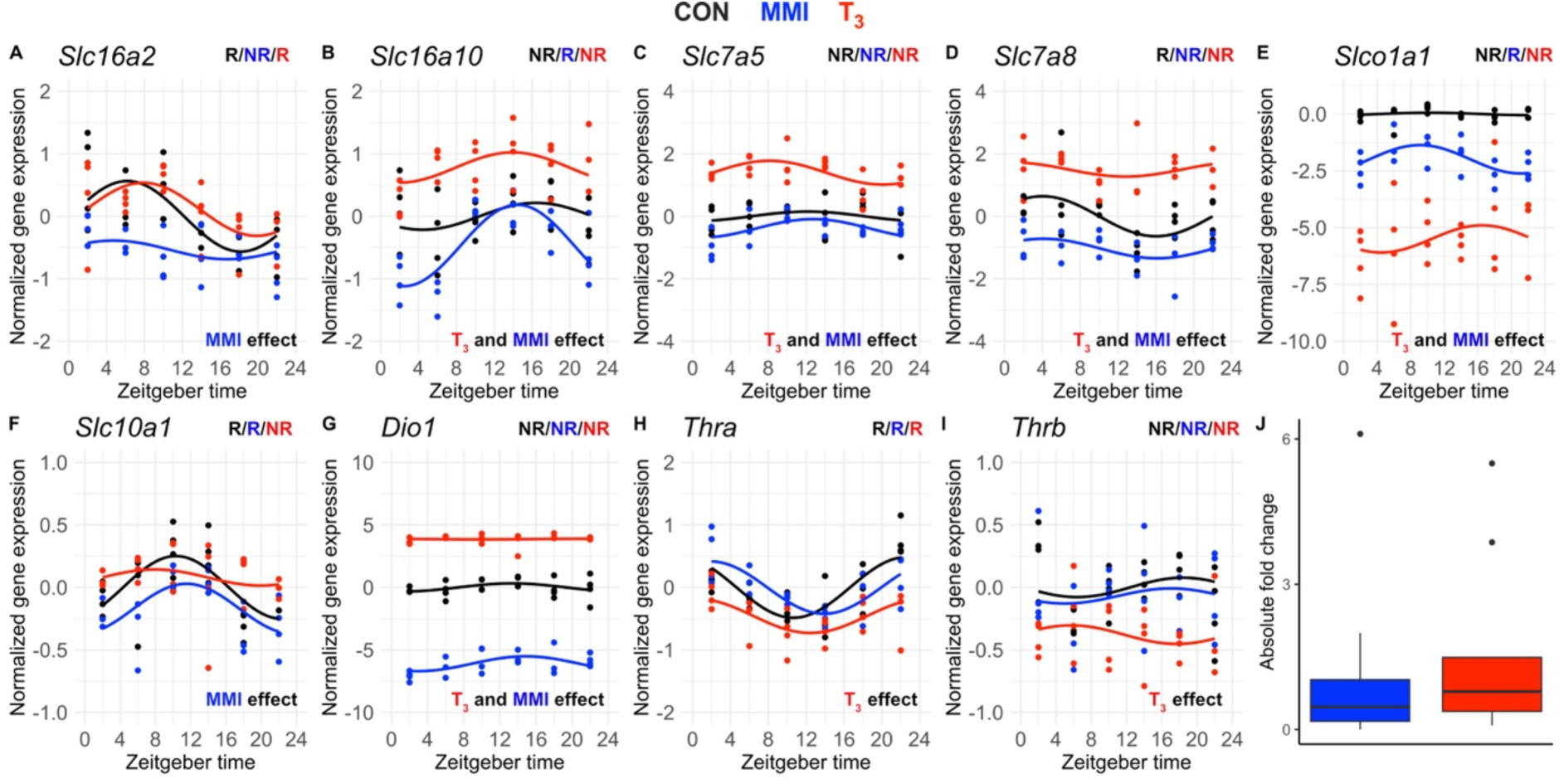
Expression of thyroid hormone regulator genes is affected by low and high TH state. A – I) Diurnal expression profiles of selected classical TH regulator genes are shown. Presence (R) or absence of significant circadian rhythmicity (NR) by JTK cycle (p value < 0.05) is depicted. J) Average absolute fold change comparisons of all TH regulators (n = 9) are shown (non-significant). Effects of low- or high- TH states were estimated using two-way ANOVA (main treatment effects, (p < 0.05). n = 3 – 4 for all ZTs and groups.

**Supplementary figure 3:**
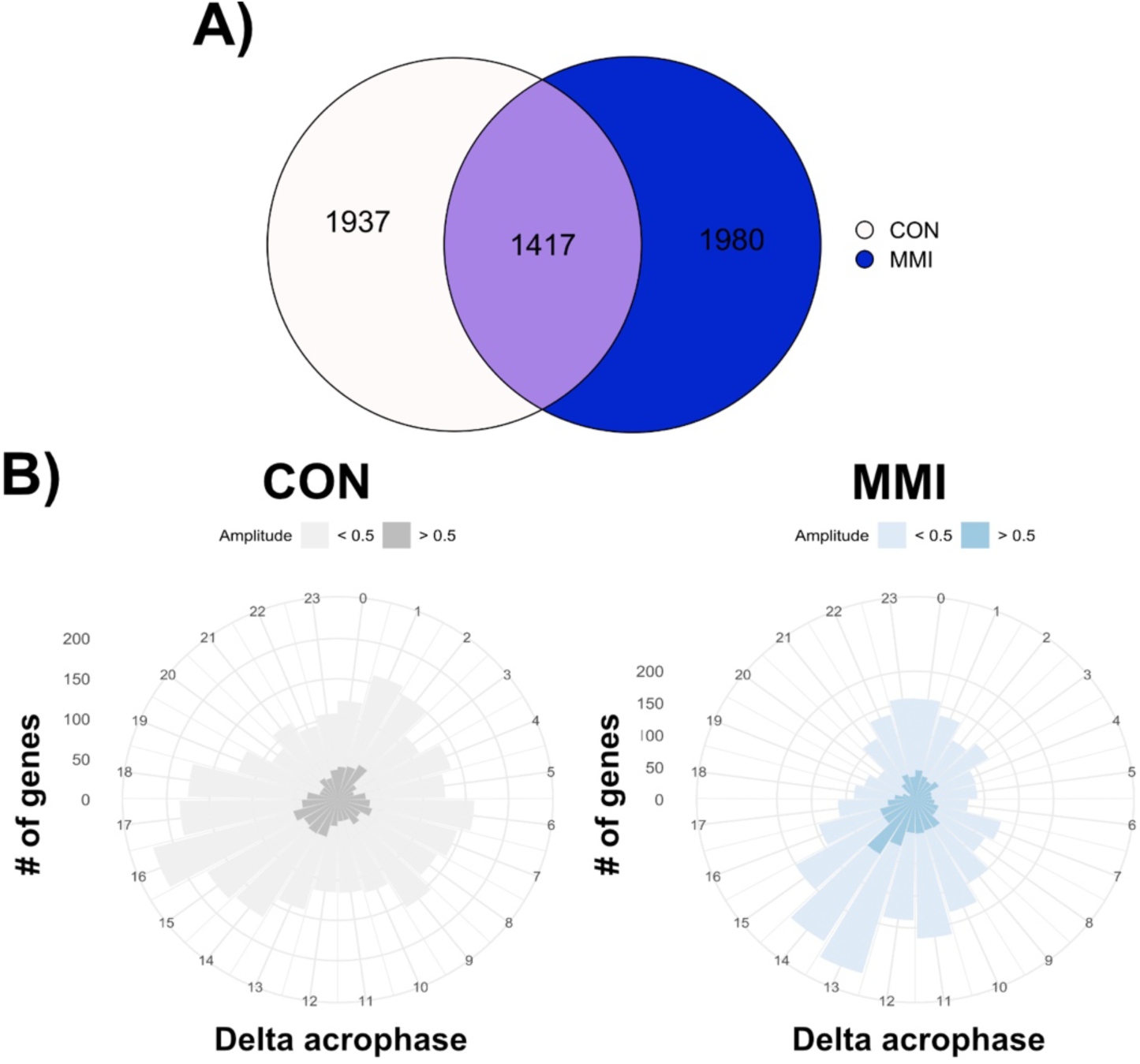
Liver diurnal transcriptome characterization in low TH conditions. A) Venn diagram showing significantly rhythmic probes identified in CON and MMI groups (JTK cycle, p < 0.05). B) Rose plots (depicting peak phase) of rhythmic genes in CON and MMI are shown.

## Notes

### Competing Interest Statement

The authors have declared no competing interest.

